# The evolution of mutation rates in the light of development and cell-lineage selection

**DOI:** 10.1101/2025.01.16.633395

**Authors:** Paco Majic, Malvika Srivastava, Justin Crocker

## Abstract

Mutation rates drive the pace and potential of evolutionary change. However, to better understand the evolutionary implications of mutation rates, there is a need to uncover the causes of their diversification. In multicellular organisms, all mutations first arise in a single cell in a developmental context. Whether a mutation enters a population’s gene pool can therefore depend on developmental events that affect the likelihood of mutant cell lineages of producing gametes. For this reason, the evolution of mutation rates in populations is governed not only by changes in the rates at which mutations occur at the molecular level, but also by changes in developmental features of organisms and how mutations impact cellular fitness during development. We present a theoretical framework that, supported by empirical data from mammals and experiments on the fruit fly, demonstrate how generational mutation rates can be shaped by changes in developmental parameters and intraorganismal selective processes even when molecular mutation rates are presumed constant. Our model highlights how the diversity of mutation rates observed across animals may be the byproduct of organismal development rather than the result of direct selection against mutator alleles. As such, development not only introduces phenotypic biases, it also shapes the rates and trajectories of genetic diversity and is thus at the core of evolutionary theory.

## Main Text

> “Seen from this point of view, life appears as a current which goes from germ to germ through the intermediation of a developed organism” - Henri Bergson, *Creative Evolution*

## Introduction

How genetic variation arises within a population shapes evolutionary trajectories (1–8). The rates of mutations in a population may affect how promptly adaptive genotypes may be encountered, while also defining how deleterious mutations may burden populations. Thus, mutation rates are closely tied to the pace and potential of evolutionary change. Although classical models presumed substitution and mutation rates to be relatively constant across the tree of life (9, 10), advances in molecular biology have progressively challenged this view (11–14). The depth and granularity at which we can now explore genetic diversity suggest that mutation rates are an evolvable trait, varying not only between clades (15–17), but also within species (18) and populations (19), underscoring their dynamic role in shaping genetic diversity.

The diversification of mutation rates can be shaped by the evolution of mutator alleles, defined as “any genomically encoded modifier of the mutation rate” (20). These alleles are typically considered to operate at the molecular scale, altering the balance between DNA damage and repair through variations in genes involved in these processes (14, 17, 20–23). Mutator alleles that promote hypermutation may be favored under strong adaptive pressures (19, 24), while those causing excessive deleterious mutations would be selected against (25, 26). Because most mutations tend to be deleterious, the mutation rates of species are thought to be dominated by the efficacy of selection in purging alleles that increase the rates of mutations (26). This is why the drift-barrier hypothesis posits that the efficiency of selection against mutator alleles depends on population size. Larger populations can better counteract genetic drift, resulting in lower mutation rates, whereas smaller populations experience weaker selection against mutators and, thus, higher mutation rates (20, 26, 27). This view suggests that mutation rates evolve primarily through changes in molecular mechanisms constrained by selection. However, mutation rates may also evolve through non-molecular causes. A relevant factor in this regard is generation time and age at reproduction, which influence the number of mutations germlines accumulate before reproduction (28–31).

Another non-molecular factor that may influence mutation rate evolution is that, in multicellular organisms, new mutations necessarily arise in a single cell in the organismal context of each parent. For a mutation to enter the population’s gene pool, it must propagate within the organism and be part of the germline, meaning developmental traits have the potential to shape the likelihood of heritable mutations. For instance, the timing of germline determination (32), and differences in gamete production between sexes or species can influence mutation rates (16, 30, 31, 33). Moreover, mutations may result in differences in cellular fitness, which can greatly influence the genetic diversity existing within organisms. In the germline context, mutations that affect the fitness of cells relative to their sisters may bias their own heritability if they occur on the lineage of cells that go from zygote to gamete (34–41). Such bias can have important evolutionary repercussions (40): it may affect the probability of fixation of variants that alter cell fitness, it allows for a “cheap” purging of deleterious variants, and it can also promote evolvability when the impact of mutations on cellular fitness and on individual fitness align. Nevertheless, cell-lineage selection tends to be overlooked in discussions of mutation rate evolution (17, 22, 30), and it is generally unaccounted for in mutation rate estimations, since whenever selection is considered there is a tendency to focus exclusively at the level of individuals - a perspective which biases causal explanations of mutation rate variation (36).

In this work, we emphasize the view that differences in population-level mutation rates across species may result from causes other than molecular changes mediated by mutator alleles. Using analytical models, data analysis and experiments in *Drosophila,* our work consolidates a view of mutation rate evolution, which demonstrates how the interplay of developmental and life-history parameters with intraorganismal selection on mutant cells can majorly affect the rate of new mutations in a population. This perspective has important conceptual implications for understanding the evolution of mutation rates. Unlike explanations like the drift-barrier hypothesis, our model highlights how developmental processes may influence generational mutation rates, positioning them as a byproduct of organismal development rather than direct selection at the level of individuals in a population. Furthermore, this model broadens the role of organismal development in population genetics: development not only introduces phenotypic biases, but it also shapes the rates and trajectories of genetic diversity, which places development at the core of evolutionary theory.

## Results

### A theoretical framework to assess the influence of developmental features and cell-lineage selection on mutation rate diversification

To distinguish between the molecular mutation rate, *μ*_mol_, and the mutation rate per generation at the population level, *μ*_pop_, we revisited the theoretical framework of Otto & Orive (35), who explored how the mutation load of a population is affected by intraorganismal selection within individuals (see Methods). Conceptually, development is modeled as a branching tree of cell lineages, with *μ*_pop_ partly determined by the number of cells carrying a mutation at locus *i* at the end of development, *M_i_* (Figure 1A). *M_i_* is a function of:

i. a molecular variable *μ*_mol_, which is the mutation probability per cell per locus per unit of developmental tempo, which can be either per cell division when mutations are of replicative nature (35), or per time unit when they result from other sources of DNA lesions (29, 42),
ii. developmental parameters such as the duration of development (*τ*), and the rate of cell proliferation (*C*), and
iii. the effect of mutations at locus *i* on cell fitness (*β_i_*), which alters the rate of cell proliferation of mutant cells relative to nonmutant cells in the developmental context. *β_i_* < 1 indicates deleterious effects that decrease cell proliferation, *β_i_* = 1 neutrality, and *β_i_* > 1 beneficial effects that increase proliferation.

**Figure 1.**
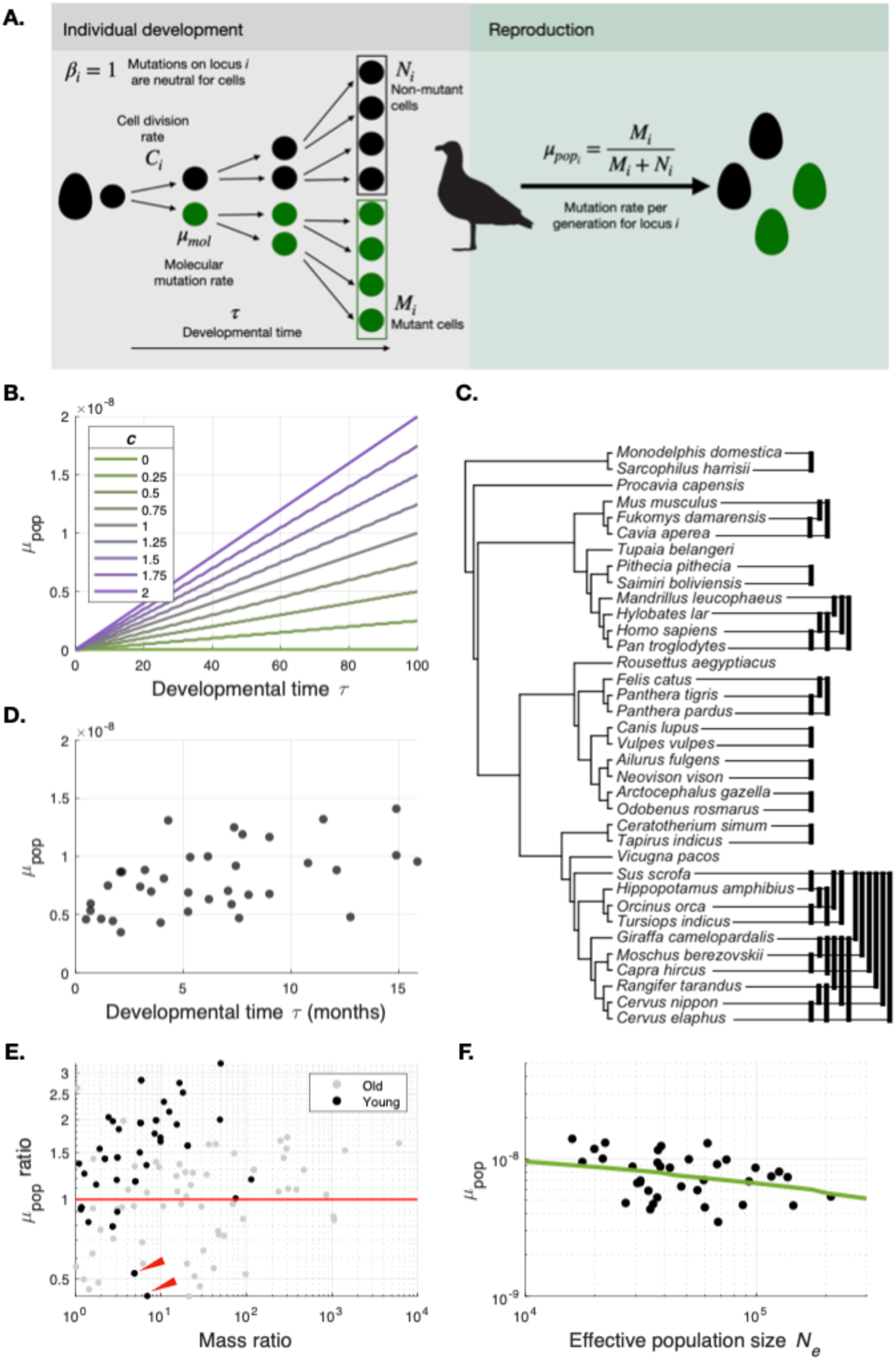
Changes in developmental parameters can causally explain diversification of mutation rates. (A) Illustrated summary of the core model consisting of an individual developmental phase and a reproductive phase. In the developmental phase a lineage starting from a single cell goes through successive cell divisions at a rate *C* over a developmental time τ. During development, mutations can occur at a rate *μ*_mol_. Cell proliferation could be affected by the fitness effect of mutations on cells *β*. For *β* = 1 mutations are neutral. At the end of development, during the reproductive phase, the fraction of germline-capable cells that carry mutations define the rate at which new alleles are added to the gene pool *μ*_pop_. (B) Population-level mutation rate *μ*_pop_ at a single locus *i* as a function of τ. Each curve represents a different value of *C.* *μ*_mol_ was held constant at a value of 1×10^-10^ mutations per cell division, approximating empirical estimates for human and mouse (91). (C) Phylogeny of mammals for which estimations for *μ*_pop_ are available (16). Black bars indicate recently diverged species pairs for which body size and *μ*_pop_ were considered. (D) *μ*_pop_ for each mammalian species in (C) as a function of their gestation time, a proxy for τ. (E) Ratio of the *μ*_pop_ of species pairs as a function of the ratio of the mass of species pairs. Black dots indicate branchings younger than 65 my, while gray dots represent older branchings. Notice how, for the vast majority of comparisons between younger branches *μ*_pop_ ratios are above 1 (red line), which means that larger clades tend to have higher mutation rates, as predicted by our model. Red arrowheads indicate the species pairs corresponding to the tiger (*Panthera tigris*) and leopard (*Panthera pardus*) and goat (*Capra hircus*) and dwarf musk deer (*Moschus berezovskii*) (F) Datapoints represent the expected relationship between the predicted *μ*_pop_ of each of the studied mammalian species and their effective population size *N_e_* based on their body mass (Methods). The green line represents the prediction of this relationship based on our developmental model for *μ*_pop_.

With an estimation of *M_i_* and assuming that germline determination is independent of mutational events, it is then possible to calculate *μ*_pop_ at Iocus *i* as the probability of sampling a mutant cell among the total pool of cells, which includes all *M_i_* mutated cells as well as all cells that completed development without mutations, *N_i_*. Such that:

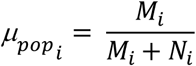

By defining *μ*_pop_ as the probability of randomly sampling mutant cells at the end of development, we are implying that this probability also encompasses the probability that sampled cells are of the germline lineage that goes from the zygote to differentiated gametes. This captures aspects of development such as how body size, which scales with the total number of cells, generally reflects germline size (43). However, our model does not consider many nuances of actual development. Development is highly structured with different cell lineages proliferating and differentiating at different tempos. Furthermore, overproliferation of cells by means of increases in *β_i_* could result in tumorigenesis, which would affect *μ*_pop_ through deleterious effects on individual fitness. However, more often than not, mutant alleles spread selectively within healthy tissues without causing tumors or deleterious effects on organismal fitness (44–50). This is not only true for somatic tissues, but also for the germline (41). Our formalization is therefore sufficient to assess how fundamental developmental features in combination with cell-lineage selection can influence the rates at which mutations enter a population’s gene pool, regardless of their effect on organismal fitness and even if the molecular mutation rate *μ*_mol_ is constant.

### Developmental parameters can influence the diversification of mutation rates

As per our model, under neutral conditions (*β_i_* = 1), evolutionary lineages that increase *τ*, also increase the expected *μ*_pop_ for locus *i* (Fig. 1B). This is also the case for shifts in *C*, but only if *μ*_mol_ is considered to track cell divisions (Fig. 1B). When considering *μ*_mol_ to track mutations per unit time, *μ*_pop_ also increases as a function of *τ*, but it is only affected by changes in *C* only if changes in the rate of cell proliferation also result in changes in *τ* (Fig. S1A). To evaluate whether these developmental parameters may explain interspecific differences in *μ*_pop_, we analyzed estimates of *μ*_pop_ (16) for a number of amniote species. Focusing on the mammalian clade, which has the most comprehensive phylogenetic sampling (Fig. 1C, Table S1) and developmental information (51), we focused on gestation time as a proxy for *τ*, as well as on the body mass as a proxy for total cell count, proportional to 2*^C^*^τ^ (52). As predicted by our model, the *μ*_pop_ of mammals positively correlated with the proxy *τ* (Fig. 1D, Pearson’s *ρ* = 0.49, *p* = 0.0025), as well as 2*^C^*^τ^ (Fig. S1B, Pearson’s *ρ* = 0.48, *p* = 0.0029).

To more directly address the question of whether developmental parameters could account for the interspecific diversification of *μ*_pop_, we compared these variables between sister clades in our phylogeny by calculating the ratio of *μ*_pop_ and body mass (Methods). We found that the majority of the comparisons between relatively recently diverged species (Fig 1C, black bars) yielded a correlation such that the larger of both species tended to have higher *μ*_pop_ (Fig. 1E, black dots). In 79% of recently diverged species pairs the larger of both species also had a higher *μ*_pop_, which was significantly different from random pairs (Fig. S1C, *p* = 0.01, one-sample permutation test). For example, the estimated *μ*_pop_ of domestic cats (*Felis catus*) is ∼2 times lower than that of tigers (*Panthera tigris*) and ∼2.5 times lower than that of leopards (*Panthera pardus*), while the *μ*_pop_ of walruses (*Odobenus rosmarus*) is ∼1.95 times that of the much smaller Antarctic fur seals (*Arctocephalus gazella*). Interestingly, this held not only for mammals but also for birds, even if the phylogenetic granularity of estimated *μ*_pop_ is lower (Fig. S2, in 74% of comparisons of related bird species pairs the larger of both species also had a higher *μ*_pop_, *p* = 0.02, one-sample permutation test). Therefore, as lineages diverge, we expect that if either lineage tends to increase in size, its mutation rate will also tend to increase. We do note that body size is also correlated with generation time, and that the correlation between generation time and *μ*_pop_ (16), as well as the influence of parental age at reproduction (53), may bias our interpretation. But, if developmental parameters affecting body mass or other correlated life history traits are indeed affecting *μ*_pop_, then the diversification of developmental parameters may offer a non-molecular explanation for observed diversification patterns of mutation rates.

As mentioned above, according to the drift-barrier hypothesis, the negative correlation existing between the generational mutation rates of species and their effective population sizes could be due to the efficacy of selection against alleles affecting *μ*_mol_ (20, 26). However, that relationship could also be the neutral byproduct of changes in developmental parameters affecting body size, rather than the selective purging of mutator alleles. This is because mammals follow a global scaling relationship between body size and population density described by Damuth’s rule (54), as well as with the effective population size (17). Considering that body mass is proportional to the total number of cells in a body, then *τ* and *C* not only affect *μ*_pop_, but also the demographic properties of a population. Therefore, simply considering the predicted effective population size for different values of *Cτ* based on this proportionality produces an inverse relationship between the mutation rate of a species and its effective population size (Fig. 1F), as would also be expected from the drift-barrier hypothesis. Thus, the differences in mutation rates observed in populations of different sizes may be the result of changes in developmental parameters that, because of ecological and energetic reasons, may also relate to changes in population sizes (55).

### The embryonic selective environment and its effect on mutation rates

So far, we have been considering the case in which mutations are selectively neutral at the cellular level. In other words, we have been considering the case where *β_I_* = 1. However, if we take into account how mutations may impact their own heritability by affecting cellular fitness, a selective regime in the embryo or body of an organism can then also affect *μ*_pop_ without changes in *μ*_mol_ (Fig. 2A). Cellular fitness could be affected by various processes, including the migration of cells, proliferation, and, in the specific case of germline cells, the determination and maintenance of the germline cell fate. Mutations on genes involved in these processes may end up under- or over-represented in the gamete pool produced by an organism (41, 56).

**Figure 2.**
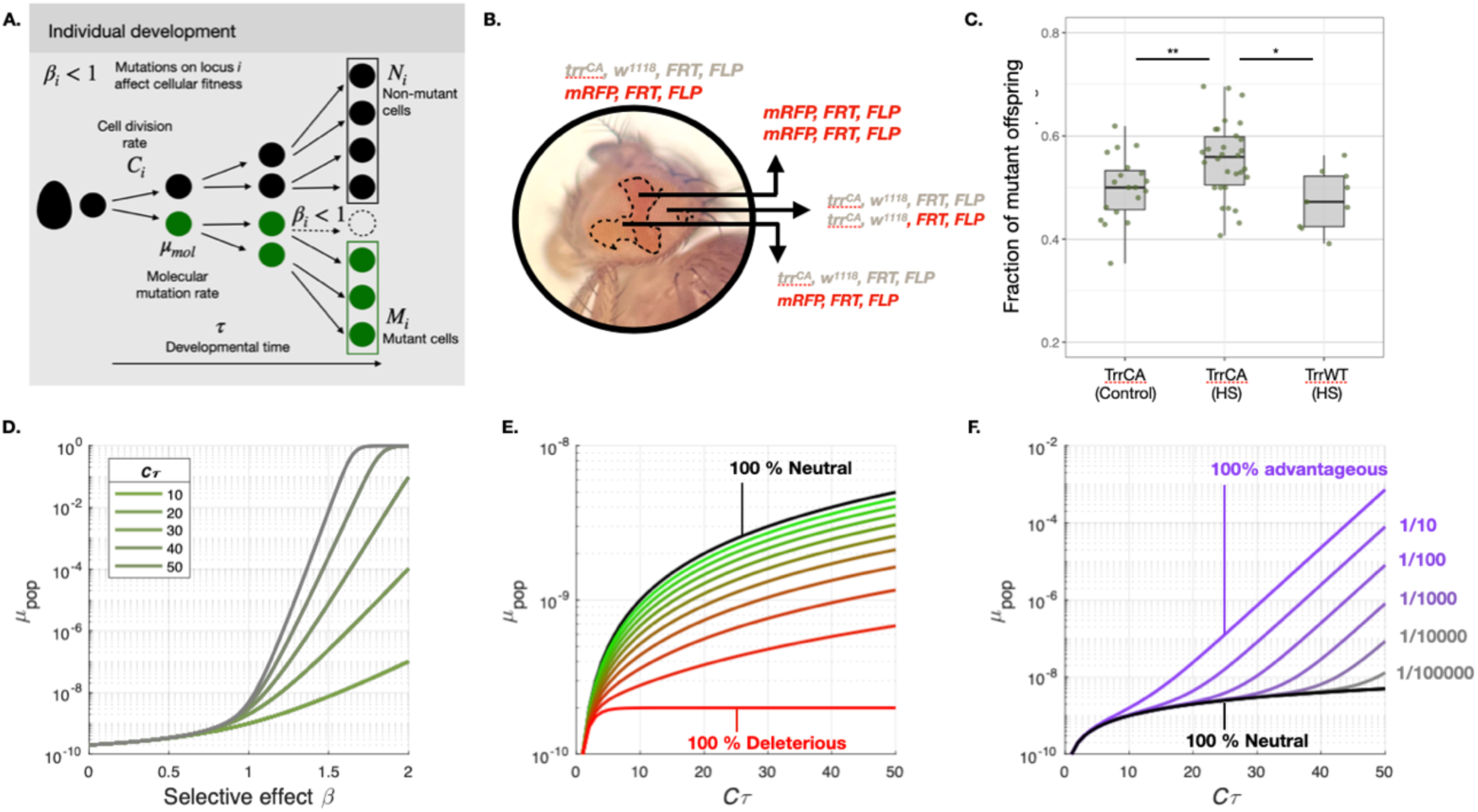
The impact of mutations on cell fitness during development can repercuss on mutation rates. (A) In the model introduced in Fig. 1A, when *β* ≠ 1, mutations can affect the total number of mutated cells at the end of development *M*. (B) Induced mitotic recombination can generate mosaic organisms with cells with different genotypes. The shown specimen corresponds to a female *Drosophila melanogaster* on which mosaicism can be easily observed in the eyes. Red, white, and orange patches represent each of the possible genotypes thanks to the *w*^1118^ allele. (C) Fraction of offspring carrying the mRFP(-) allele from crossing a female of interest with a *w*^1118^ male. TrrCA refers to females of the *trr*^CA^,*w*^1118^ genotype, while TrrWT refers to the *w*^1118^ genotype that is wild-type for *trr*. “HS” refers to females that developed from embryos that were heat-shocked, while “Control” females developed from embryos that were not heat-shocked. Each point represents the fraction of offspring from a different mother. * indicates *p* < 0.05, ** indicates *p* < 0.005 using a Wilcoxon test. (D) Population-level mutation rate *μ*_pop_ as a function of the effect of mutations on the fitness of cells *β*. Each curve represents a different value of developmental time τ with a constant value for *C =* 1. (E) Population-level mutation rate *μ*_pop_ as a function of developmental parameters *C*τ. Each curve represents a different value proportion of neutral or deleterious mutations. (F) Population-level mutation rate *μ*_pop_ as a function of developmental parameters *C*τ. Each curve represents a different proportion of mutations that are advantageous for the cell (*β* = 1.5) in the context of an otherwise fully neutral sequence.

To explore how mutations affecting *β_i_* as in our model may impact their own heritability, we considered the *Drosophila melanogaster* gene *trithorax-related (trr)*, which influences cell proliferation (57). We generated mutant clones by inducing mitotic recombination through the FLP/FRT technique (58) on flies that carried both a sub-functional allele of *trr* (TrrCA,(59)) on an unmarked chromosome and a wild-type version of the gene (TrrWT) on a chromosome marked with a *red fluorescent protein, mRFP* (see Methods). As a result, within a single embryo, some cells are homozygotic for TrrCA, homozygotic for TrrWT, or heterozygotic with a copy of each allele (Fig. 2B). Because an impaired *trr* activity increases the rates of cell proliferation (57), we then expected that *β_trrCA/trrCA_* > *β_trrWT/trrCA_* > *β_trrWT/trrWT_* and, therefore, that the TrrCA allele would be overrepresented in the germline and thus have a higher heritability than TrrWT. Consistent with this hypothesis, when mosaicism was induced early in development, a median of ∼55% of offspring carried the mutant allele, a fraction that was significantly higher than the cases when mosaicism was not induced on the TrrCA/TrrWT heterozygotes or when it was induced in the lines with two copies of TrrWT (Fig. 2C). This experiment illustrates how a mutation inducing a greater proliferation of cells that carry it can increase its own heritability, representing a case of what Otto and Hastings referred to as “mitotic drive” (40).

In our model, varying the proliferative effect *β_i_* of a mutation on a cell can drastically change the effective *μ*_pop_ even when molecular mutation rates are constant, such that higher values of *β_i_* imply higher *μ*_pop_ (Fig. 2D). A relevant insight from this model is that even mild consequences for cellular proliferation may have major consequences for the rate at which new mutations enter the gene pool of populations. For example, in the case of *C*τ = 20 (representing 2^20^ cells, roughly the number of ovarian follicles an infant is born with (60) or half of the number of germ cells in the ovary of cows by the third trimester of pregnancy (61)), a mere 1% increase (*β_I_* = 1.01) or decrease (*β_i_* = 0.99) in cell fitness may result in a ∼7% change in *μ*_pop_ (Fig. 2D).

The effect of *β_i_* on *μ*_pop_ also depends on the developmental parameters *C* and τ (Fig. 2D). Higher values of *C*τ tend to produce larger differences in *μ_pop_* for large*β_i_*. On the other hand, when *β_i_* is small, *C*τ does not have a strong effect on the mutation rate. We note that when *β_i_* ≠ 1, changes in *C* and τ, also affect *μ*_pop_ when *μ*_mol_ is considered as mutations per unit of time, rather than per cell division (Fig. S3A). This contrasts with the aforementioned neutral case, where changes in *C* did not change *μ*_pop_ for constant τ (Fig. S1A).

Our model, therefore, suggests that: i) the selective consequences of mutations at the cellular level may influence generational mutation rates, and ii) the magnitude of this influence depends on developmental parameters. Specifically, our model predicts that organisms of larger size and slower development should be more efficient in purging mutations that are deleterious for cells and more prone to facilitate the fixation of mutations that are beneficial to cells (Fig. 2D). This shows how differences in mutation rates between species may be enhanced by developmental differences, and it highlights how the relevance of cell-lineage selection as a “cheap” way to filter deleterious mutations (40) may be evolve by shifting developmental parameters.

### The interplay between cellular distribution of fitness effects and developmental parameters

The effect of mutations on cellular fitness is expected to vary across genes, depending on their involvement on cellular processes, and even across loci within genes, depending on their relevance for gene function. For example, in our *Drosophila* case study, despite carrying a mutation in the gene *white* (*w*) (Methods), the TrrWT allele did not lead to differences in heritability in our FLP/FRT experiments (Fig. 2C). This suggest that even though mutations on *trr* can influence *β_i_*, mutations on *w* do not, which implies that *trr* and *w* likely contribute differentially to *μ*_pop_.

To explore how differences in *β_i_* at different loci affects genome-wide mutation rate estimations, we adjusted our developmental model such that the overall genomic mutation rate would be proportional to the fraction of sites that have either deleterious (*F*_*β* <1_), neutral (*F*_*β*=1_) or advantageous (*F*_*β* >1_) fitness effects on cells (Methods). Moving from a case in which mutations on all sites are neutral for cellular fitness (*F*_*β*=1_ = 1,), to a case in which all mutations are strongly deleterious (*F*_*β*=0_ = 1), there is a steady drop in the expected *μ*_pop_, tending towards *μ*_mol_ as most mutations become deleterious (Fig. 2E). Considering cases in which most mutations are either neutral or deleterious for cellular fitness would be in accordance with what is expected (25). In contrast, scenarios in which many mutations tend to increase cellular fitness are presumably rare. However, our model does suggest that even if infrequent, mutations with strong beneficial effects on the fitness of cells can drastically influence the genome-wide estimations of *μ*_pop_. Depending on the developmental parameters *C*τ, this is true if mutations that increase cellular fitness are 1 in 1000 (*F*_*β* >1_ = 0.001), or even 1 in a million (*F*_*β* >1_ = 1*x*10^−6^) (Fig.2F). Notably, variations in the ratios of *F*_*β*=1_, *F*_*β* >1_ and *F*_*β* < 1_ also vary the relationship between *μ*_pop_ and developmental parameters (Fig. 1E,F).

Furthermore, because *β_i_* affects mutated cells by altering their proliferative capacity, the number of genome-wide mutations capable of increasing or decreasing cellular fitness may depend on the rate of cell proliferation of wild-type cells. In other words, *F*_*β*=1_, *F*_*β* >1_ and *F*_*β* <1_ may depend on baseline *C,* such that for greater *C* there should be a smaller number of mutations for which *β_i_* > 1. Such would represent a case of diminishing returns epistasis, according to which the fitness effect of a mutation is milder when it occurs on a fit genotype than when it occurs on an unfit genotype (62–64). Epistatic effects of this kind could explain why there is a negative correlation between our estimates of *C* for a subset of mammalian species (Methods) and *μ*_pop_ (Fig. S3B, Pearson’s *ρ* = −0.38, *p* = 0.044), since greater *F*_*β* >1_ imply greater *μ*_pop_ (Fig. 2F). This would mean that whenever species evolve a greater size through increases in *C*, the cellular distribution of fitness effect could change as well, thus affecting *μ_pop_*.

Therefore, the genome-wide distribution of cellular fitness effects of mutations interacts with and may be defined by the developmental background of an organism to jointly influence *μ*pop. A factor that needs consideration in any interpretation of differences in mutation rates between species using genomic data.

### Selection in the mature germline stem cell niche

In addition to embryonic development, the lifetime of organisms could also shape *μ*_pop_. This could take place, as mentioned earlier, because longer generations imply a greater accumulation of new mutations before reproduction (30, 65, 66). Here, we argue that this could also be because of the dynamic competition between germline stem cells of different genotypes. During an individual’s lifetime, germline stem cell niches can undergo a cellular turnover. In mice, for example, spermatogenic stem cells sometimes cease to be stem cells and are replaced by neighboring stem cells, keeping the number of stem cells relatively constant (67). Although the repopulation of these cleared niches by neighboring cells could be a stochastic process (68), mutations that affect the rate of stemness loss (either through death or differentiation) might eventually create a mosaic of spermatic stem cells with an overall higher fitness (41, 56). Thus, a selective cell turnover might impact the repertoire of mutations that a single individual can introduce into a population’s gene pool.

To incorporate the effect of selection during the lifespan of an organism, we built on our developmental model described above and included a maturation and reproductive phase (Fig. 3A, Methods). In this adapted version, at the end of development, there is a final and constant pool of stem cells that can produce gametes, *M_t=_*_0_ of which are mutated. These cells can undergo a turnover, either because of a death-birth dynamic in the stem cell niche or because of differentiation. This turnover of cells in the stem cell niche occurs at a rate *d* over an arbitrary unit of time *t* subdividing the total generational time *l*, and a new selective variable *β*_*m*_ which describes the fitness effect of mutation on cells related to stem cell turnover during the lifetime of an organism. In this model, in the case when there is no turnover of cells during the lifetime of an individual (*d* = 0) or when mutations are neutral (*β*_*m*_ = 1), *μ*_*pop*_ only changes due to the mutational accumulation of new mutations. Whenever there is a turnover of cells and mutations do affect the proliferative capacities of cells, a positive effect of mutations on the fitness of cells increases the overall *μ*_*pop*_ while deleterious mutations decrease it closer to *μ*_*mol*_ (Fig. 3B). This effect is more pronounced for individuals that take longer to reproduce (Fig. 3C).

**Figure 3.**
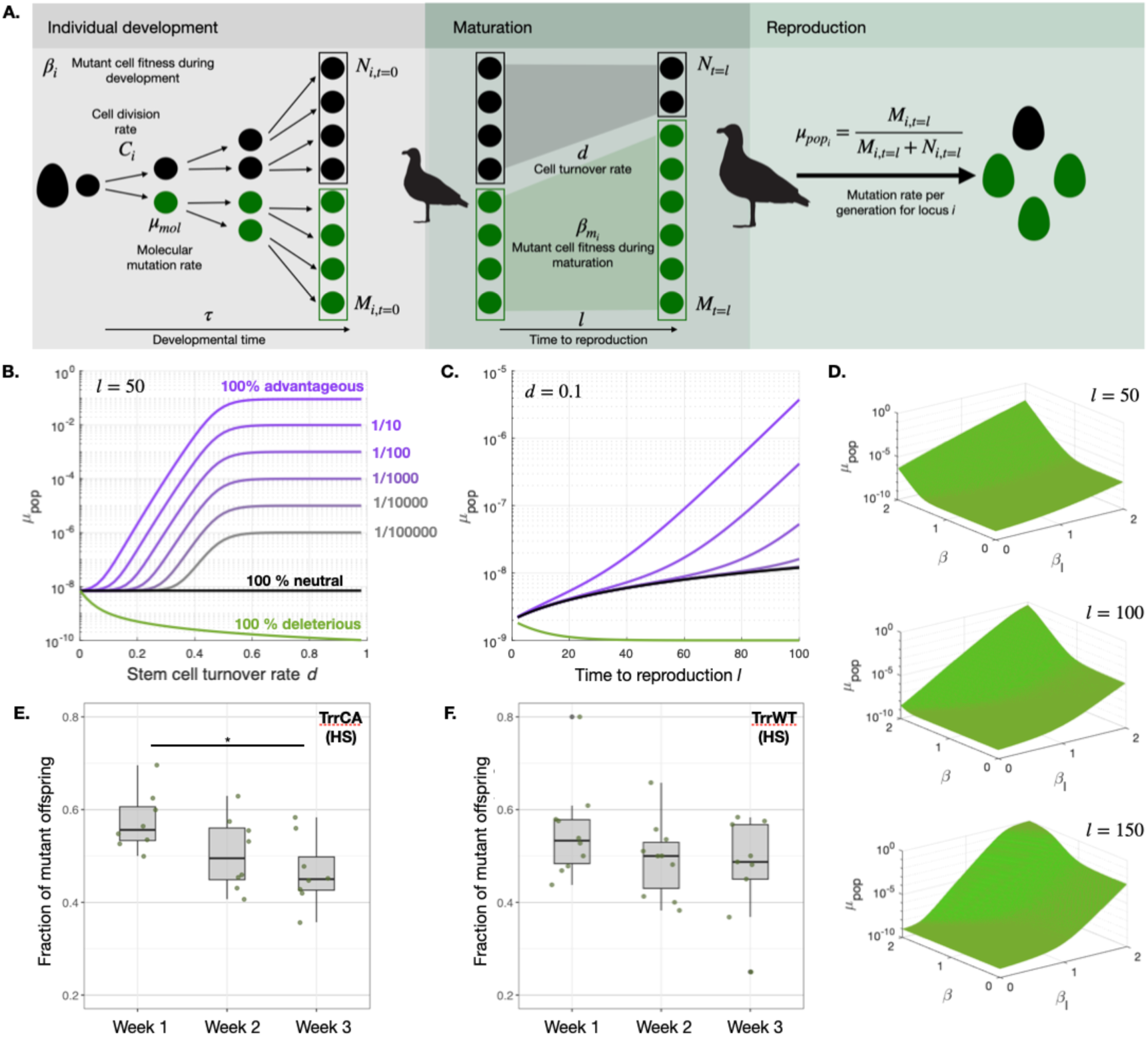
Selection within the germline stem cell niche can also impact mutation rates over an organism’s lifetime. (A) Illustrated summary of the model integrating a maturation phase before reproduction. Adding this phase incorporates parameters such as the time to reproduction of an organism *l*, a turnover rate of germline stem cells *d*, and a cellular fitness variable for the stem cell niche *β_l_*. (B) Population-level mutation rate *μ*_pop_ as a function of the rate of stem cell turnover *d* for a constant generation time of *l = 50*. Each curve represents a different proportion of mutations that are advantageous for the cell (*β_l_* = 1.5) in the context of an otherwise fully neutral sequence (gray to purple lines), a fully neutral case (black) or a fully deleterious case (green). (C) Population-level mutation rate *μ*_pop_ as a function of generation time *l* for a constant rate of stem cell turnover *d* = 0.1. The different lines follow the same color code as in (B). (D) *μ*_pop_ as a function of cellular fitness variable for the stem cell niche *β_l_* for different values of *l* and the fitness effect of mutation on cells during development *β*. (E and F) Fraction of offspring carrying the mRFP(-) allele from the same set of mothers reproducing at the beginning of weeks one, two or three post-hatching. TrrCA refers to females of the *trr*^CA^,*w*^1118^ genotype, while TrrWT refers to the *w*^1118^ genotype that is wild-type for *trr*. “HS” refers to females that developed from embryos that were heat-shocked. * indicates *p* < 0.05.

The intraorganismal selective process over the lifespan of a mature organism may interplay with the intraorganismal selective process taking place earlier during development to shape an effective *μ*_*pop*_ (Fig. 3D). For example, mutations on genes involved in basic cellular functions would likely have fitness consequences for cells at any stage of an individual’s life, thus affecting cell proliferation during development, but also the rates of germline stem cell turnover (*β* > 1, *β_m_* > 1). On the other hand, there may be genes that are irrelevant for cellular fitness in the germline stem cell niche that may be important during development (*β* ≠ 1, *β_m_* = 1) and vice versa (*β* = 1, *β_m_* ≠ 1). The latter would be the case of mutations on the *chimno* mutants in *Drosophila*, which have been reported to evict wild-type cells from the stem cell niche of the testis (38). It is also possible that the selective regime during development and maturity point in opposite directions, which is what we observed in the case of the TrrCA mutation presented above: even though mosaic females introduced in Fig. 2C produced a progeny that was biased in favor of the *trr* mutant allele right after pupation, as those same females aged, the biased shifted in favor of TrrWT (Fig. 3E), suggesting that for this allele *β* > 1 and *β_m_* < 1. On the other hand, as expected, TrrWT retained a *β* = *β_m_* = 1 through the development and lifespan of each female fly (Fig. 3F).

These results reinforce the notion that the estimation of global genome-wide mutation rates may lead to biased interpretations of the molecular rates because of the fitness consequences of mutations on different loci. More generally, our results also support that the overall expected *μ*_*pop*_ is likely the composite outcome of the selective effect of mutations at the cellular level during development and maturity, as well as the life history traits of the organism.

## Discussion

Mutations arise in organismal contexts that vary across individuals and species. As our results demonstrate, these contexts significantly influence how genetic variation is produced and incorporated into gene pools. Therefore, differences in population-level mutation rates may not solely result from changes in molecular mutation rates or changes in generation time, but could arise from evolutionary shifts in developmental parameters. These include the rate of cell proliferation, developmental duration, germline differentiation modes, germline dynamics during maturity, and—crucially—selective dynamics at the cellular level.

This perspective implies that it is not essential to invoke direct selection on mutation rates to account for the observed diversity of mutation rates across multicellular lineages. This is not to say that selection at the level of individuals plays no role in modulating mutation rates. Molecular rates might indeed evolve, for example, to mitigate the burden of deleterious mutations, particularly somatic mutations that can cause cancer in the case of larger and longer-lived species (65, 69–71). However, because developmental parameters alone can produce substantial shifts in mutation rates, as species evolve different morphologies and life history strategies, their rates of genetic diversification can also diverge as a secondary outcome that may or may not be neutral.

Our results illustrate how the diversification of mutation rates can arise as a consequence of changes in developmental parameters associated with body size evolution. This is supported both by our theoretical model and by a general pattern in which the larger of two closely related species tends to exhibit a higher generational mutation rate. However, this trend is not universal and can be influenced- or even reversed-by other factors. Although body size typically correlates with germline size (43), deviations from this relationship, such as disproportionate germline cell numbers or developmental heterochronies in the gonads, can alter expected generational mutation rates, as suggested to explain higher rates in chimpanzees relative to humans (72). Additionally, developmental changes tied to body size may interact in complex ways with intraorganismal selection. For instance, in the leopard-tiger and goat-dwarf musk deer species pairs (Fig. 1C), the larger species shows a lower estimated generational mutation rate (red arrowheads in Fig. 1E, Table S1). We speculate that this misalignment reflects a smaller fraction of mutations capable of further increasing cellular proliferation (*F*_*β* >1_) in species with an inherently higher proliferative capacity of wild-type cells. We infer tigers and goats to have a higher rate of cell proliferation per unit time (greater *C*) than leopards and dwarf musk deer, respectively. This is evidenced by the fact that tigers, despite reaching larger birth sizes than leopards, have similar developmental times, while dwarf musk deer, though significantly smaller than goats, possess markedly longer developmental times (Table S1).

### The biological causes of mutation rate diversification

The diversification of developmental parameters not only provides a basis for hypotheses about species-specific shifts in shifts in mutation rates, but also offers potential explanations for well-established patterns of mutation rate diversification. Our results suggest that the prevalent negative correlation between generational mutation rates and population size (16, 17, 20, 26), is more parsimoniously explained by developmental variations accompanying lineage divergence than by recurrent selective adjustments of molecular mutation rates in response to demographic changes, as proposed by the drift barrier hypothesis (26). Given that ecological drivers may influence body size evolution (55) and that developmental parameters that increase body size also increase generational mutation rates, the co-adjustment of population size and developmental properties (or correlated life history traits) could underlie the diversification of generational mutation rates. Although the drift barrier hypothesis is supported by various studies that show a correlation between mutation rates and effective population size across clades (17), we are here suggesting that, at least for multicellular organisms, there are other biological properties that may explain the same trend, or at least reinforce it. This is consistent with our findings and supported by evidence that diversification in mutation spectra, often attributed to molecular changes, primarily occurs between distantly related lineages (73) and may also be influenced by environmental factors (74) or cell cycle dynamics (23).

The developmental influence on generational mutation rates also offers alternative explanations to other patterns of mutation rate variability that have been attributed to changes in molecular mutation rates. An example is the differential accumulation of mutations during different stages of mouse development, which has been interpreted to be the result of a stage-specificity of DNA repair systems (75), but that could be explained by a stage-specificity of any of the developmental parameters we explored. Another example is the negative correlation between the generation times of species and their estimated yearly mutation rates (76). A proposed explanation aligning with the drift barrier hypothesis is that gamete mutation rate modifiers are constrained by the product of population size and generation time by selection at the individual level (65). Our intra-organismal selection model predicts similar trends, albeit through a different mechanism: longer-lived species allow more efficient elimination of variants that are deleterious the cellular level and a greater retention of beneficial ones due to extended selective dynamics. Thus, if most mutations tend to negatively affect cellular fitness, which, as discussed above, may be the case for larger organisms that also have longer generation times, intraorganismal selection would produce lower estimations of yearly mutation rates for species with longer generation times.

Developmental parameters and intraorganismal selection shape not only interspecific differences in mutation rates, but may also help explain variation within species. A well-established pattern is the higher mutation rates in males compared to females in the germlines of mammals and birds, a bias largely absent in reptiles and fish, where reproductive biology is more similar between sexes (16). The male-bias in birds and mammals has been attributed to a greater cell division number during the lifetime of males (30), but also to imbalances in the DNA repair mechanisms between the sexes (77). But sex-specific biases could also be caused by intraorganismal selection (40, 41, 56). The selective expansions of mutant lineages in the male germline may exacerbate the differences in generational mutation rates, including the paternal age effect (78), as has been reported from clinical cases (41, 79, 80). In terms of our model, male-specific expansions are facilitated by a greater stem cell turnover over the lifetime of organisms, with females having *d* ∼ 0 and males *d* > 0 (Fig. 3B). Interestingly, if intraorganismal selection is at least partially driving the differences between male and female mutation rates, we expect not only a quantitative bias, but also a qualitative bias: whereas mutations inherited from females would be enriched in mutations that are deleterious to cells and would likely get purged if that mutation arose in the male germline, mutations inherited from fathers are likely enriched in effects increasing cell fitness.

Although our argument broadly aligns with the idea that non-molecular causes may underlie the diversification of generational mutation rates (28, 29, 33), we do not rule out the evolutionary relevance of molecular mechanisms. Alterations to baseline molecular mutation rates are well-documented (81) and likely coexist with intra-organismal selection and developmental influences (36, 82). Disentangling whether a particular molecular, developmental or life history parameter has driven the diversification of generational mutation rates between species may be challenging, as many relevant factors tend to be correlated. But ultimately, the causes of mutation rate diversification are likely multifaceted. In this context, we propose an expansion of the definition of “mutator alleles” (20) beyond molecular mechanisms that affect DNA repair and damage. Instead, mutator alleles may also expand to genes that influence developmental processes, life history traits, or organismal physiology.

### Considerations about the place of development in evolution

The role of development in evolution is increasingly recognized, but much of the discussion has focused on how developmental systems constrain or drive phenotypic evolution (83–86). Our findings emphasize an additional role: by influencing mutation rates, development can shape the evolutionary pace and direction at both phenotypic and genotypic scales.

In the conceptual framework of genotype space (87), populations could, in principle, explore the space equally in all directions (Fig. 4A). However, mutational biases—such as the well-documented transition versus transversion bias—skew this exploration (Fig. 4B), affecting the spectrum of adaptive substitutions available to populations (4). Beyond molecular biases, developmental processes can further constrain or promote specific directions of exploration (Fig. 4C) and influence the breadth of genotype space sampled within a generation (Fig. 4D). This has quantitative and qualitative implications for the genetic diversity of populations. Quantitively, populations of larger organisms like rheas, for example, can explore a broader portion of genotype space compared to those of smaller organisms like gulls, which have a considerably smaller generational mutation rate (16). While qualitatively, this broader exploration may occasionally introduce deleterious variants, many of these—particularly those disrupting basic cellular functions or germline potential—are more efficiently pruned in larger species due to their developmental and intraorganismal selective dynamics. Thus, organisms are not merely vehicles for genes, as major evolutionary paradigms suggest (88). Instead, development actively participates in shaping genotypic changes arising in the gene pools of populations and, by extension, the evolutionary trajectories of species.

**Figure 4.**
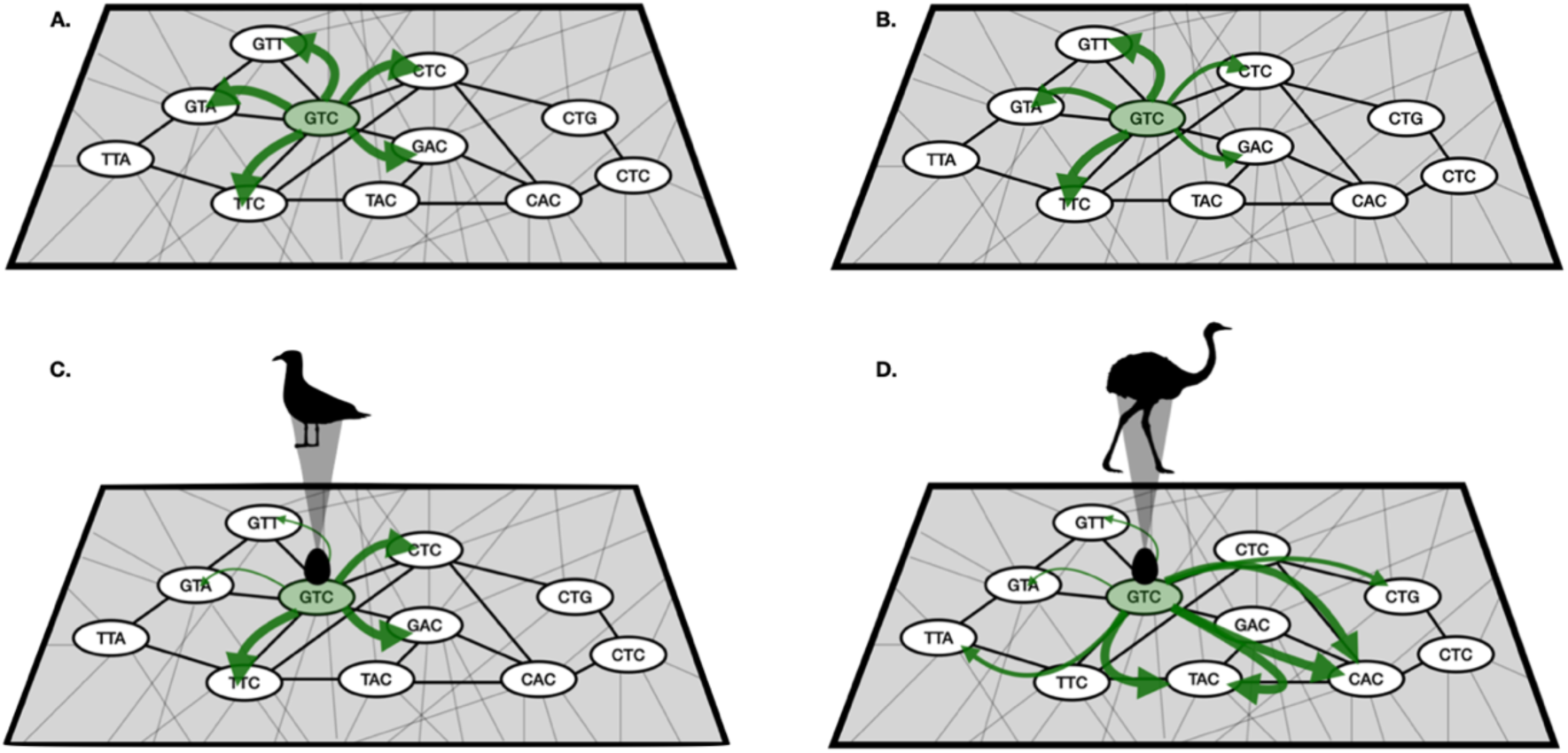
How development can influence the trajectories and pace of genotypic evolution. In a theoretical model of evolution, it could be considered that given a starting genotype, GTC, all mutational steps are equally likely (A). However, if molecular realism is considered there are certain mutations that are more likely than others, as represented by arrows of different thickness going to CTC and GAC. (B). Furthermore, as we argue in this work, development can act as a selective filter for mutations which increases or decreases the likelihood of mutational steps depending on how a mutation affects the cell lineage it belongs to (C). Developmental and life-history differences can also impact how far mutational steps can take an organism. As per our model, larger organisms should be capable of more broadly exploring genotypic spaces within a generation (D).

Overall, by highlighting the interplay between development and selection at the cellular level, we provide a broader framework for understanding the diversification of mutation rates. Taking this framework into account, we suggest that future studies exploring the diversification of mutation rates should account for the developmental contexts in which mutations arise in each species and the ecological interactions between cells of different genotypes.

## Materials and Methods

### Developmental model for estimation of *μ*_*pop*_

We define the generational mutation rate *μ*_*pop*_ as the probability of sampling a mutant cell among all cells in an organism that are capable of producing gametes at the time of reproduction of an individual. To calculate this probability, it is necessary to first calculate *M_i_*, which represents the total number of cells that are expected to carry a mutation at a given locus *i*. We adapted a model from Otto & Orive (35), wherein *M_i_* is a function of the molecular mutation rate *μ*_*mol*_, the total developmental time τ, the rate of cell divisions during development *C*, and a factor *β*_i_, which represents the effect of a mutation on the fitness of a cell carrying a mutation on locus *i*. At developmental time *x*, one would expect 2^*x*^(1 − *μ*_*mol*_)^*x*−1^ cells to not carry any mutations. Each of those cells can mutate with a probability given by *μ*_mol_ and the mutated lineage will then undergo *β_i_*(*C*τ *-x*) cell divisions more until the conclusion of development. Thus, the number of mutant cells resulting from mutations at time *x* is:

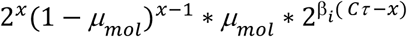

Summing over all *x* gives us *M_i_* at the conclusion of development:

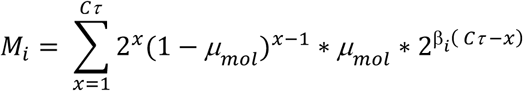

For estimating *μ*_pop_ it is also necessary to calculate the expected number of cells that do not undergo mutations at locus *i* through the course of development i.e., *N*_i_. This can be calculated as:

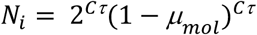

The total number of cells at the end of development is (*M*_*i*_ + *N*_*i*_) and thus 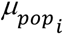 approximates the probability of sampling a cell mutated at locus *i* from all cells in the pool of gametes, such that

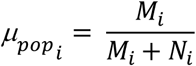

The calculations shown above correspond to those considering *μ*_*mol*_ to track cell divisions. In the cases in which mutation rates are taken to track time rather than cell divisions, the calculations for *M_i_* and *N_i_* were modified such that,

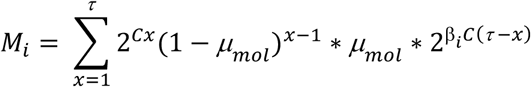

And,

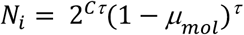

To estimate *μ*_pop_ across the entire length of a genome, as a first order approximation, we assumed that no cell obtains mutations at more than one locus during the entire course of development. In other words, we assume *μ*_*mol*_^2^ ≈ 0. This assumption would hold for very small *μ*_mol_. Using this simplification, we can estimate *μ*_pop_ across the entire length of a genome by averaging *μ*_*pop*_across all sites with different values of *β*_i_. In our model, we considered *F*_*β*i=*f*_ to be the fraction of the total sites in a genome that have a *β*_i=_ *f,* giving:

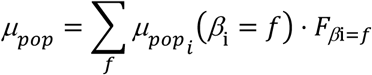

### Expansion of the model to account for maturation and time to reproduction

To consider the role of lifetime of an organism in the calculation of *μ*_*pop*,_we considered the pool of cells resulting from the developmental process to be constant in number. Immediately at the end of development, there are *M_t=_*_0_ cells in the germline stem cell niche that are mutated and *N_t=_*_0_ that are not. During an organism lifetime *l,* germline stem cells can undergo a turnover (due to differentiation or death) at a rate *d* over an arbitrary unit of time *t*. The rate at which mutant cells replace lost cells during this turnover is given by a selective variable *β*_*m*_. As in the case of the developmental selective parameter *β*, when *β*_*m*_ < 1 mutations are deleterious, when *β*_*m*_ = 1 mutations are neutral and when *β*_*m*_ > 1 mutations are advantageous to cells. In our model, during the lifetime of an organism before reproduction, it is possible for the total pool of mutated cells to decrease either due to reverse mutations (which would occur at a low rate, *M*_*t*–1_*μ*_*mol*_) or loss due to death or differentiation (*dM*_*t*–1_). Furthermore, mutant cells can increase in proportion whenever non-mutant cells in the stem cell niche mutate (*N*_*t*–1_*μ*_*mol*_) or whenever mutant cells proliferate to replenish the stem cell niche after the loss of sibling cells. The number of mutated cells at time *t* during the lifetime of an organism can thus be calculated as,

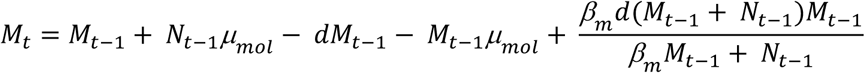

And the number of non-mutant cells would be given by

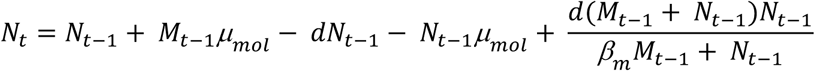

### Comparison of *μ*_*pop*_ for different mammalian species

To corroborate the predictions of our model we considered estimations of the *μ*_*pop*_ of each of 36 species of mammals. We considered the *μ*_*pop*_ of each species to be the mean of the number of de novo mutations per base pair across different numbers of parent-offspring trios (16). We collected the other biological parameters used, namely the body mass of each species and the gestation time, from the Encyclopedia of Life (51). The phylogenetic relationships we considered were based on recent phylogenetic assessments of mammals (89), while the estimated divergence times were collected from various sources listed in Table S1. To estimate *C*, we considered gestation times as a proxy for τ and the mass of neonates as a proxy for 2^*C*τ^ for those species for which this information was available in the Encyclopedia of Life.

To consider how the mutation rates of recently diverged species have evolved in relation to developmental parameters, we compared the ratios of the body mass for pairs of species with the ratios of mutations rates measured by Bergeron et al (16). The pairs we used for our comparisons consisted of pairs of the two most closely related species (Fig. 1C). In cases like those of *Sus scrofa* that have more than one species that are equally closely related, we considered all possible species pairs (Fig. 1C, black bars). For these comparisons, we considered the ratio of the mutation rate or body size of the smaller of both species against that of the larger one. Thus, the ratio of body sizes should be greater than 1 for all comparisons. If the ratio of mutation rates is also greater than one, body size and mutation rates are aligned, while if it is smaller than one, the smaller species have a greater mutation rate. To test the significance of our result, we compared the alignment of the *μ*_*pop*_ and body size ratios (the percentage of comparisons for which *μ*_*pop*_ ratios were greater than 1) with the expectation from 1000 random permutations of *μ*_*pop*_. Pairs of species were considered to be recently diverged if their common ancestor dates back to less than 65 mya (black bars in Fig. 1C, older pairs are not shown in the phylogeny but we used them for our comparisons, Fig. 1E, gray point). The same analysis was carried out for the birds species for which mutation rates were available (16), but no phylogenetic aging was assigned since all extant birds diverged after the Cretaceous-Tertiary boundary of 65 my.

### Estimation of effective population size (*N*_*e*_) and the relationship between *μ*_*pop*_ and *N*_*e*_

Organisms comprising of 2^*Cτ*^ cells, have a mass *m* = *m*_*cell*_ 2^*Cτ*^, where *m*_*cell*_ = 10^−15^*g* is the mass of a single cell (90). We used this mass estimate to compute the effective population size *N*_*e*_. This was done by using the scaling of effective population size and organism size (adult dry weight) that was reported by Lynch et al. (17) for organisms across the tree of life. The fitted log– log regression for the data resulted in the line: y = 6.717 − 0.199x. The above mass estimate was also used to calculate the mutation rate *μ*_*pop*_using our equation. Finally, we also compared our predicted curve with mammalian weight and mutation rate data (see section above). Setting *μ*_*mol*_ to 10^−9.74^gave the best agreement with the empirical data.

### Flippase assay to assess cell competition in fly germlines

We developed heat shock-induced mitotic clones in fruit flies following a standard protocol (58). Briefly, a fly line (# 31418 from the Bloomington Drosophila Stock Center) containing a red fluorescent protein (mRFP), a flippase-encoding gene induced by a heat-shock promoter (hsFLP) and a FRT site for the binding of flippases was crossed with another line carrying a single-amino acid substitution in the *trithorax-related* gene (*trr*) and the *w*^1118^ allele for the gene white. We thus created a *trr*^CA^,*w*^1118^,FRT,hsFLP and a *w*^1118^,FRT,hsFLP lines. For our experiments we crossed these lines with the parental mRFP,FRT,hsFLP lines and we exposed the heterozygotic embryos collected during a window of four hours to a heat shock of 37°C for one hour.

We collected mosaic virgin females that developed from these heat-shocked embryos as well as virgin females from embryos that were not heat-shocked (control). Each of these virgins were individually paired to two *w*^1118^ males and we then counted the fraction of offspring that were positive for red fluorescence, which are the ones that inherited the mRFP+ allele and not the mRFP-(*trr*^CA^,*w*^1118^) allele. For the same set of pairs, we evaluated the fraction of mRFP+ pupae from offspring produced during the first week immediately after mothers were collected, as well as during the second and third weeks.

## Acknowledgments

We are grateful to Lautaro Gandara, Anna-Lena Vigil, David Porubsky and Cecilia Padilla Iglesias for their useful feedback on our manuscript.

## Supplementary materials

**Table S1.**
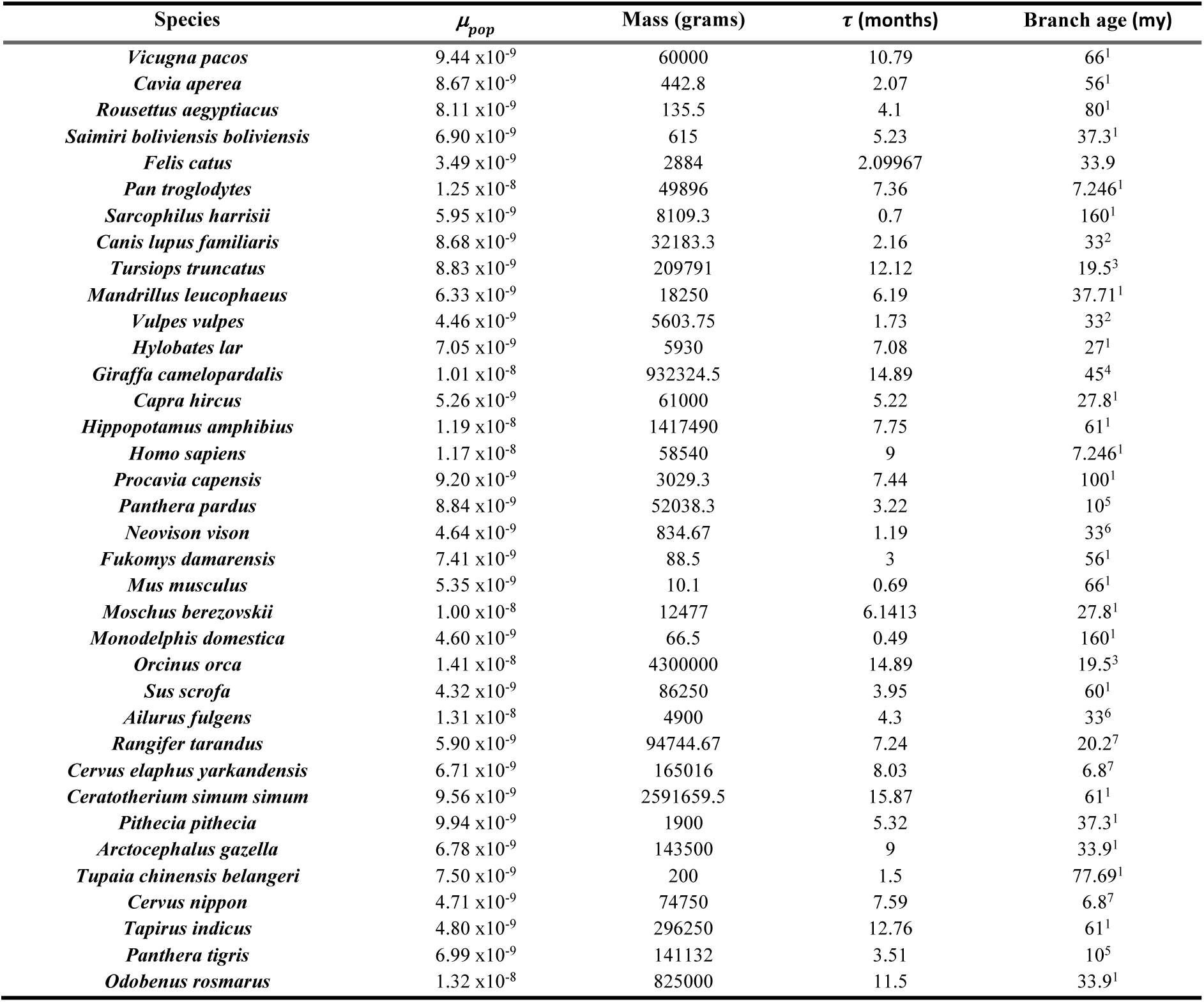
Empirical parameters for 36 mammalian species. Bergeron et al (2023) estimated the number of de novo mutations in parent-offspring trios of each of the studied species. With those estimations we calculated *μ*_pop_ as the average of all replicates reported. The mass and τ (gestation time) values were collected from the Encyclopedia of Life (Parr et al 2014). Branch ages representing the estimated divergence time from the closest relative in the phylogeny (Fig. 1C) were gathered from several sources indicated by the superindeces in the Branch Age column: ^1^(89), ^2^(92) ^3^(93) ^4^(94) ^5^(95) ^6^(96) ^7^(97).

| Species | $\mu_{pop}$ | Mass (grams) | $\tau$ (months) | Branch age (my) |
| --- | --- | --- | --- | --- |
| *Vicugna pacos* | 9.44 x10<sup>-9</sup> | 60000 | 10.79 | 66<sup>1</sup> |
| *Cavia aperea* | 8.67 x10<sup>-9</sup> | 442.8 | 2.07 | 56<sup>1</sup> |
| *Rousettus aegyptiacus* | 8.11 x10<sup>-9</sup> | 135.5 | 4.1 | 80<sup>1</sup> |
| *Saimiri boliviensis boliviensis* | 6.90 x10<sup>-9</sup> | 615 | 5.23 | 37.3<sup>1</sup> |
| *Felis catus* | 3.49 x10<sup>-9</sup> | 2884 | 2.09967 | 33.9 |
| *Pan troglodytes* | 1.25 x10<sup>-8</sup> | 49896 | 7.36 | 7.246<sup>1</sup> |
| *Sarcophilus harrisii* | 5.95 x10<sup>-9</sup> | 8109.3 | 0.7 | 160<sup>1</sup> |
| *Canis lupus familiaris* | 8.68 x10<sup>-9</sup> | 32183.3 | 2.16 | 33<sup>2</sup> |
| *Tursiops truncatus* | 8.83 x10<sup>-9</sup> | 209791 | 12.12 | 19.5<sup>3</sup> |
| *Mandrillus leucophaeus* | 6.33 x10<sup>-9</sup> | 18250 | 6.19 | 37.71<sup>1</sup> |
| *Vulpes vulpes* | 4.46 x10<sup>-9</sup> | 5603.75 | 1.73 | 33<sup>2</sup> |
| *Hylobates lar* | 7.05 x10<sup>-9</sup> | 5930 | 7.08 | 27<sup>1</sup> |
| *Giraffa camelopardalis* | 1.01 x10<sup>-8</sup> | 932324.5 | 14.89 | 45<sup>4</sup> |
| *Capra hircus* | 5.26 x10<sup>-9</sup> | 61000 | 5.22 | 27.8<sup>1</sup> |
| *Hippopotamus amphibius* | 1.19 x10<sup>-8</sup> | 1417490 | 7.75 | 61<sup>1</sup> |
| *Homo sapiens* | 1.17 x10<sup>-8</sup> | 58540 | 9 | 7.246<sup>1</sup> |
| *Procavia capensis* | 9.20 x10<sup>-9</sup> | 3029.3 | 7.44 | 100<sup>1</sup> |
| *Panthera pardus* | 8.84 x10<sup>-9</sup> | 52038.3 | 3.22 | 10<sup>5</sup> |
| *Neovison vison* | 4.64 x10<sup>-9</sup> | 834.67 | 1.19 | 33<sup>6</sup> |
| *Fukomys damarensis* | 7.41 x10<sup>-9</sup> | 88.5 | 3 | 56<sup>1</sup> |
| *Mus musculus* | 5.35 x10<sup>-9</sup> | 10.1 | 0.69 | 66<sup>1</sup> |
| *Moschus berezovskii* | 1.00 x10<sup>-8</sup> | 12477 | 6.1413 | 27.8<sup>1</sup> |
| *Monodelphis domestica* | 4.60 x10<sup>-9</sup> | 66.5 | 0.49 | 160<sup>1</sup> |
| *Orcinus orca* | 1.41 x10<sup>-8</sup> | 4300000 | 14.89 | 19.5<sup>3</sup> |
| *Sus scrofa* | 4.32 x10<sup>-9</sup> | 86250 | 3.95 | 60<sup>1</sup> |
| *Ailurus fulgens* | 1.31 x10<sup>-8</sup> | 4900 | 4.3 | 33<sup>6</sup> |
| *Rangifer tarandus* | 5.90 x10<sup>-9</sup> | 94744.67 | 7.24 | 20.2<sup>7</sup> |
| *Cervus elaphus yarkandensis* | 6.71 x10<sup>-9</sup> | 165016 | 8.03 | 6.8<sup>7</sup> |
| *Ceratotherium simum simum* | 9.56 x10<sup>-9</sup> | 2591659.5 | 15.87 | 61<sup>1</sup> |
| *Pithecia pithecia* | 9.94 x10<sup>-9</sup> | 1900 | 5.32 | 37.3<sup>1</sup> |
| *Arctocepalus gazella* | 6.78 x10<sup>-9</sup> | 143500 | 9 | 33.9<sup>1</sup> |
| *Tupaia chinensis belangeri* | 7.50 x10<sup>-9</sup> | 200 | 1.5 | 77.69<sup>1</sup> |
| *Cervus nippon* | 4.71 x10<sup>-9</sup> | 74750 | 7.59 | 6.8<sup>7</sup> |
| *Tapirus indicus* | 4.80 x10<sup>-9</sup> | 296250 | 12.76 | 61<sup>1</sup> |
| *Panthera tigris* | 6.99 x10<sup>-9</sup> | 141132 | 3.51 | 10<sup>5</sup> |
| *Odobenus rosmarus* | 1.32 x10<sup>-8</sup> | 825000 | 11.5 | 33.9<sup>1</sup> |

**Figure S1.**
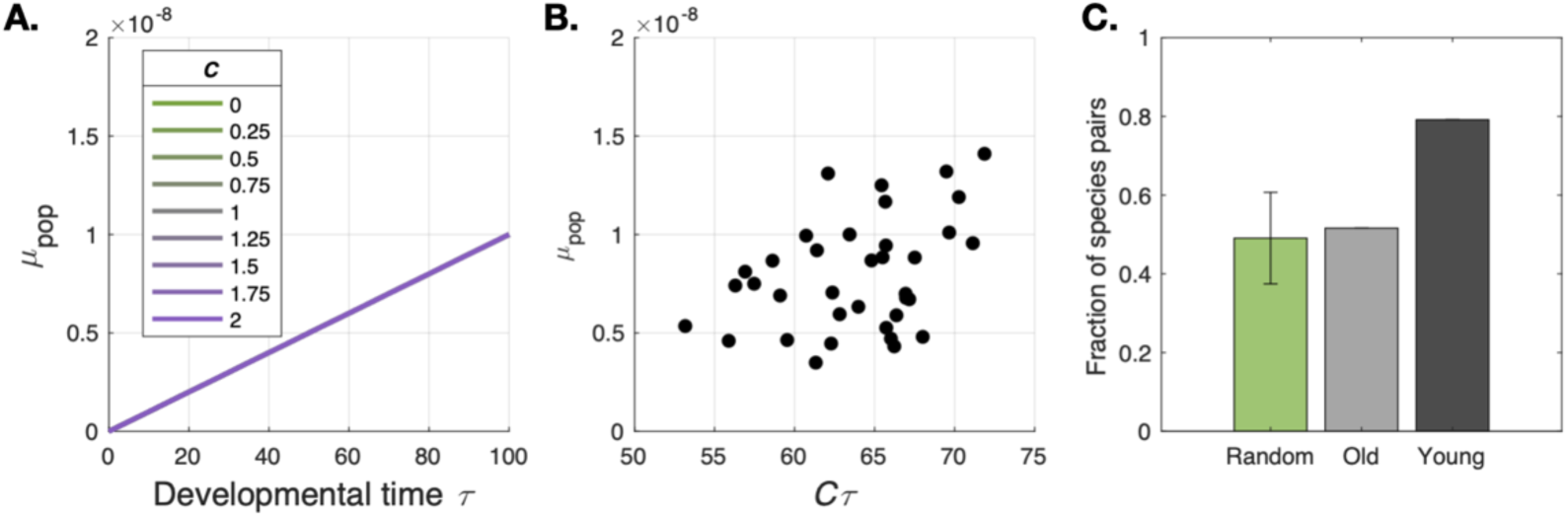
(A) Population-level mutation rate *μ*_pop_ at a single locus *i* as a function of τ. *μ*_mol_ was held constant at a value of 1×10^-10^ mutations per developmental time unit. Each curve represents a different value of *C*, but they are all collapsed on the same line. (B) *μ*_pop_ for each mammalian species in Fig. 1C as a function of *C*τ, which we estimated as the log2 of the number of cells of an organism, measured as the ratio between body mass and the mass of a single cell (Methods). (C) Barplot showing the fraction of species pairs for which the larger of both species also a higher *μ*_pop_. The green bar shows the mean fraction of 1000 random permutations of the values with error bars indicating the standard deviation, the gray bar indicates the fraction for species pairs that branched more than 65 mya, while the black bar shows the fraction for recently diverged species pairs.

**Figure S2.**
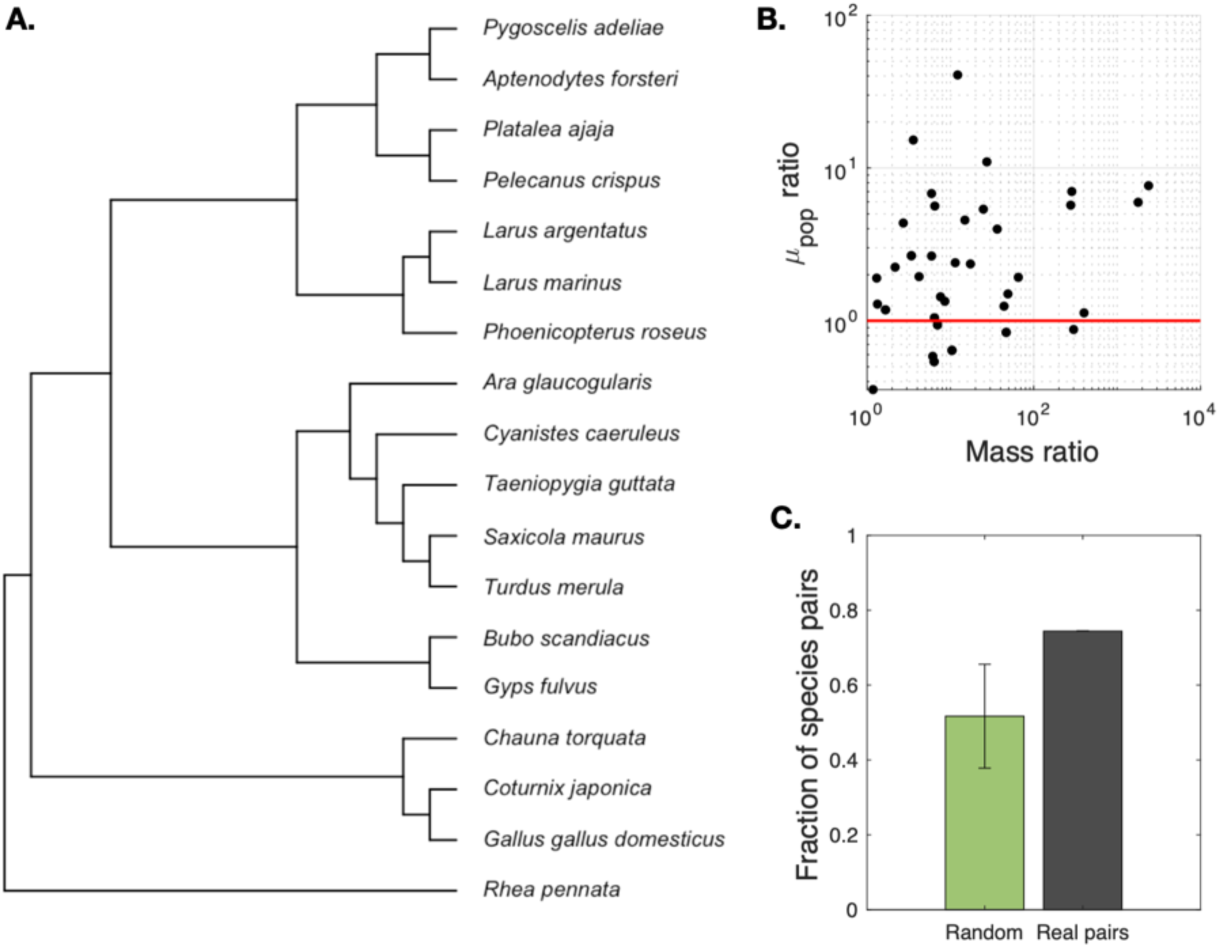
(A) Phylogeny of birds for which estimations for *μ*_pop_ are available (16). (B) Ratio of the *μ*_pop_ of species pairs as a function of the ratio of the mass of species pairs. Notice how, for the vast majority of comparisons between younger branches *μ*_pop_ ratios are above 1 (red line), which means that larger clades tend to have higher mutation rates, as predicted by our model. (C) Barplot showing the fraction of species pairs for which the larger of both species also a higher *μ*_pop_. The green bar shows the mean fraction of 1000 random permutations of the values with error bars indicating the standard deviation, while the black bar shows the fraction for all considered empirical species pairs.

**Figure S3.**
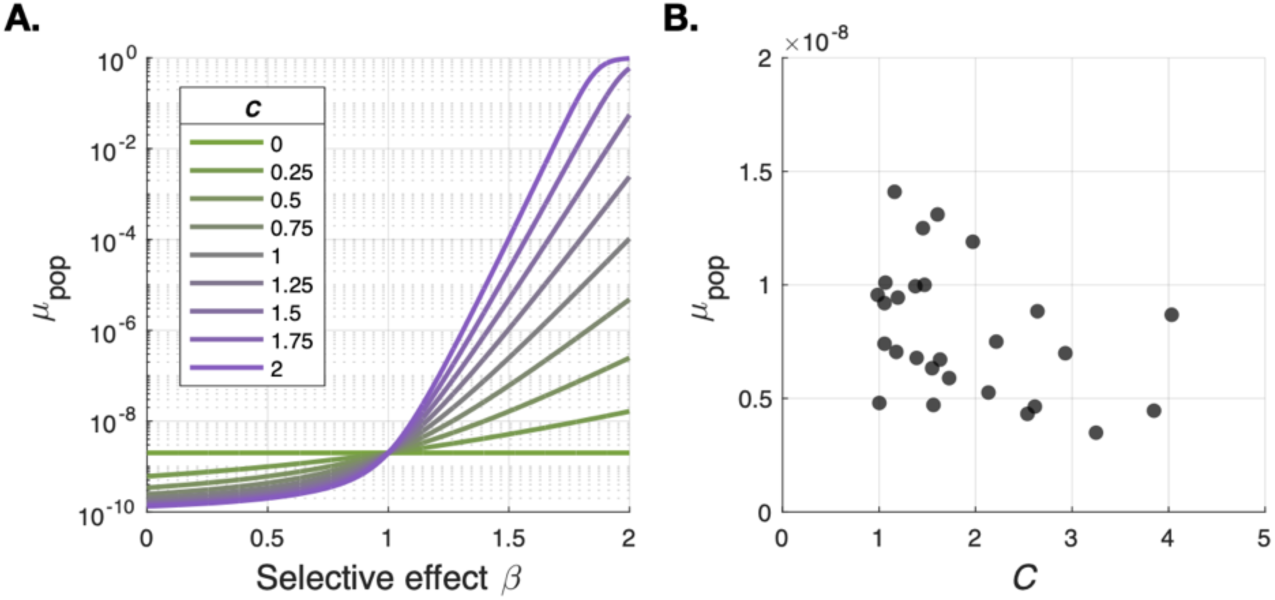
(A) Population-level mutation rate *μ*_pop_ at a single locus *i* as a function of *β*_i_ for τ = 20. *μ*_mol_ was held constant at a value of 1×10^-10^ mutations per developmental time unit. Each curve represents a different value of *C*. (B) *μ*_pop_ for each mammalian species in Table S1 for which neonate mass was available as a function of an estimated *C* (Methods).

## Notes

### Competing Interest Statement

The authors have declared no competing interest.

### Summary of Updates

Better contextualisation in the introduction and the discussion. Additional analysis are presented supporting our results.

